# Drivers of grass endophyte communities in prairies of the Pacific Northwest, USA

**DOI:** 10.1101/2020.02.23.953489

**Authors:** Graham Bailes, Dan Thomas, Scott D. Bridgham, Bitty A. Roy

## Abstract

Prairies of the Pacific Northwest are highly threatened, with only ∼2% of historic land area remaining. The combined risk of global climate change and land use change make these prairies a high conservation priority. However, little attention has been paid to the microbiota of these systems, including the hyper diverse fungi that live asymptomatically in their leaves, the endophytes. Using culture-free, full-community DNA sequencing, we investigated the diversity, composition, and structure of full fungal foliar endophyte and ecological guild communities in two native, cool-season bunchgrasses along a climate gradient. We quantified the relative importance of host, host fitness, environment, and spatial structuring in microbial community structure. We found markedly different communities between the southern and central-northern sites, suggesting a potential dispersal limitation in the Klamath Mountains. We also found that each host species was home to distinct fungal communities. Climate was the strongest predictor of endophyte community, while fitness (e.g., plant size, reproductive status, density) was less important for community structure. For both host species, seasonality contributed strongly to the variation we observed. At the ecological guild level, saprotrophs tended to decline with latitude, whereas symbiotrophs and pathotrophs both tended to increase with latitude. Our results suggest that climate change will have large consequences for these diverse fungal communities.

## Introduction

Prairies in the Pacific Northwest are critically endangered with 98% or more loss (Noss et al. 1995, Christy and Alverson 2011) as a result of habitat destruction from farming and urbanization (Noss et al. 1995), invasive species (Pfeifer-Meister et al. 2012, Lindh 2018), infilling with trees (Peter and Harrington 2014) and lack of fire (Pendergrass 1995, Clark and Wilson 2001, Hamman et al. 2011). While the plants of these prairies have been relatively well studied, their fungi have received little attention, despite playing major ecological roles as pathogens, symbionts, and decomposers. Furthermore, a large part of the biodiversity of these prairies are likely to be endophytic fungi, which are “hidden” inside the leaves of plants (Blackwell and Vega 2018). Traditionally, endophytic fungi have been described as those that live inside the host but cause no symptoms (Wilson 1995) but NGS and culturing studies frequently uncover many latent pathogens and saprotrophs from asymptomatic leaves. Here we use the term endophyte to describe all fungi living within asymptomatic plant leaves, that is, defined by habitat and regardless of their ecological role (Hardoim et al. 2008, Porras-Alfaro and Bayman 2011, Wemheuer et al. 2019). While grasses have been extensively studied from the perspective of clavicipitaceous endophyte fungal associations (‘C-endophyte’) (Rodriguez et al. 2009), which are restricted to grasses and are sometimes mutualistic (Faeth 2002, Rodriguez et al. 2009), relatively little is known about the non-clavicipitaceous (‘NC-endophyte’) diversity of native grass leaves from natural populations, but see (Higgins et al. 2014) for a tropical example, and Nissinen et. al for a study in Finland (2019).

The Pacific Northwest is predicted to experience changing climate patterns, including increases in temperature, precipitation seasonality and consecutive dry days (NOAA NCEI/CICS-NC). Paired with habitat fragmentation, there is real uncertainty over how microbial communities will respond. Here, we examine foliar endophytic diversity of two core grass species across a climate gradient using culture-free next generation sequencing (NGS), to disentangle how environment, spatial structure, and host drive community assembly, composition, and structure. Specifically: Do foliar fungal endophyte communities vary across distance and/or latitude? What factors influence endophyte community and fungal ecological guild structuring? Are the same patterns we observe in our full communities also present for different ecological guilds?

While foliar fungal endophytes are sometimes assumed to be buffered from environmental constraints such as drought (Thomas et al. 2016), dispersal often depends on environmental factors, especially temperature (Harvell et al. 2002, Roy et al. 2004) and precipitation (Huber and Gillespie 1992). Given the general paradigm of increasing fungal richness as latitude decreases (Arnold and Lutzoni 2007), we predicted that endophyte species richness will also tend to increase as latitude decreases. Further, we expected to see community structuring among plant host, ecoregion, and site, suggesting host specificity and non-random community assembly. We attempted to understand community structure in the context of community assembly, and these major filtering processes: Dispersal limitation (as a function of spatial distance), environmental filtering (local climate and elevation), and biotic interactions (host species and host fitness). We hypothesized that while many factors influence community composition, the climate would play the largest role in determining community composition because of the necessity of precipitation in both fungal sporulation and germination, and the consequences of temperature on survival. Finally, we examined how these filters affected the fungal ecological guilds (pathogens, saprotrophs, symbiotrophs).

## Methods

### Study location

This study was conducted along a 680 km latitudinal gradient in western Oregon and Washington, spanning three Ecoregions (Klamath Mountain, Willamette Valley, and Puget Trough-Georgia Basin) (Fig. S1). The climate from this region is Mediterranean, characterized by warm dry summers, and cool wet winters (Koppen-Geiger climate classification). Temperatures tend to become more extreme toward the south, while precipitation trends don’t track with latitude. However, both temperature and precipitation seasonality tend to increase as latitude approaches the equator (Fig. S2, Table 1).

**Table 1.**
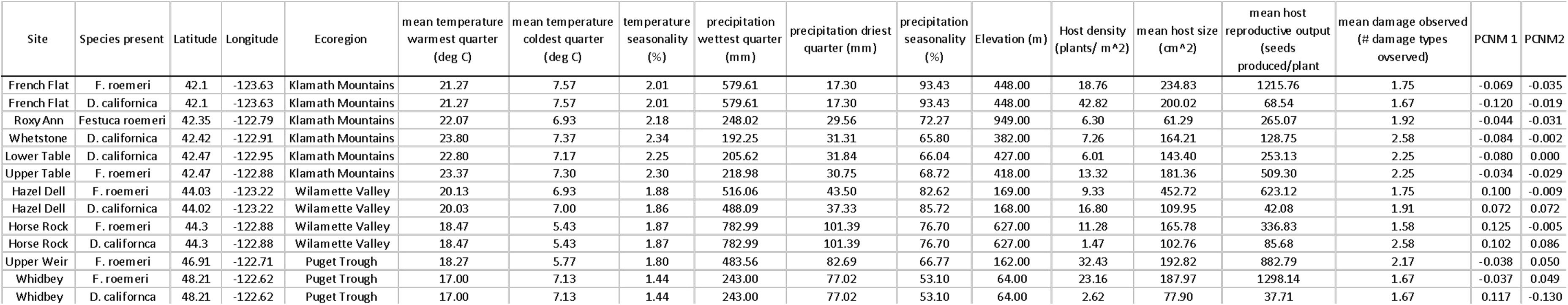

We examined endophyte communities within two native, perennial cool-season C3 bunchgrasses: *Danthonia californica* Bolander and *Festuca roemeri* (Pavlick) E. B. Alexeev. These grasses are relatively long-lived, able to survive up to several decades, and are among those species considered to be a historic cornerstone of Pacific Northwest prairies (Christy and Alverson 2011, Stanley et al. 2011). *D. californica* is widespread, with a natural range extending from southern California into British Columbia, and east from the Pacific Ocean toward the Rocky Mountains (Darris and Gonzalves 2019). *F. roemeri* is distributed from central California into British Columbia, growing only on the west side of the Cascade and Sierra Nevada mountain ranges (Darris et al. 2012). For each host species, we selected at least two sites within each ecoregion (Fig. S1). Host populations were used if they were located within remnant prairies (in which no restorative seeding had been done) and contained an adequate presence of target individuals (200+ within a 30 by 2m transect). We further strove to maximize variation in elevation, precipitation and temperature among sites to delineate a near-continuous climate gradient. In all, we sampled six populations of *D. californica*, and seven of *F. roemeri*.

### Sample collection and preparation

Five asymptomatic leaves were collected from each of 12 adult (flowering) plants from each population. These collections occurred during the summer of 2015, at peak host biomass. While we preferentially sampled for reproductive individuals in an attempt to control for plant age (which may confound endophyte load (Clay and Schardl 2002)), reproductive plants were randomly selected approximately every 2m along a 30 m transect. To model host fitness, we collected demographic data on each host individual including size, reproductive output; these fitness traits were included in the partitioning of variation analyses as “host effects”. Each host was scored for the presence of pathogen and herbivore damage types, but the leaves analyzed for endophytes showed no obvious damage. Density was measured by counting the number of plants/m^2^ along our transect.

Samples were stored on ice while in the field, transported to the laboratory to be processed within 48 hrs, at which time leaves were surface sterilized and frozen at −20 °C until DNA extraction. Surface sterilization: leaves were immersed in 70% ethanol for 1 min, then transferred to a 5% sodium hypochlorite solution for 1 min, and finally transferred to 70% ethanol for an additional minute. The leaves were then rinsed in sterilized DI water, and blotted dry. Using a flame-sterilized scalpel and tweezers, 0.5 g of leaf tissue was transferred into sterilized 2ml flat-bottomed tubes containing approximately 0.3g of sterile zirconium beads. We followed a modified protocol from the Qiagen DNeasy 96 plant kit (Qiagen, Valencia, CA, USA). After adding lysis buffer, each sample was homogenized for two cycles of 30 seconds at 3450 oscillations/minute, using a Biospec Mini Beadbeater-8 (Biospec Products, Bartlesville, OK, USA). The remaining extraction protocol followed the manufacturer’s instructions. DNA was stored at −20 °C until we constructed a metabarcode sequencing library of the internal transcribed spacer (ITS) region within the rRNA gene. Libraries were constructed as described in Thomas et al. (2019), using a duel-indexed split-barcode system. Triplicate PCR reactions were carried out in 20 µl volumes. The reagents included 4 µl genomic DNA, 0.8 µl MgCl_2_ (25 mM), 0.6 µl of each 10mM primer, 4 µl Milli-Q Ultrapure Water (MilliporeSigma, Darmstadt, Germany), and 10 µl 2x PCR Super Master Mix (100 U/ml of Taq DNA polymerase, 0.5mM dNTPs, 4mM MgCl2, stabilizers and dye) (Bimake, Houston, Texas, USA). Reactions were assembled on ice in a class II biological safety cabinet (Labconco^©^, Kansas City, MO, USA) to reduce non-specific amplification and contamination. PCR conditions included an initial five min of denaturation at 95 °C; 35 cycles of: 1 min at 95 °C, 1 min at 55 °C, 1 min at 72 °C; final elongation of 10 min at 72 °C. triplicate PCR products were pooled together and cleaned using Mag-bind^©^ Rxn PurePlus (Omega Bio-Tec^©^, Norcross, GA, USA) beads. The University of Oregon Genomics and Cell Characterization Core Facility (Institute of Molecular biology, University of Oregon, Eugene, OR, USA) completed the following Illumina preparation before samples were submitted for sequencing. Samples were normalized and pooled with those from another study (Thomas et al. (2019) to a final concentration of 7.013 ng/µl (258 samples). Fragment analysis (Advanced Analytical, Ankeny, Iowa, USA) was used to quantify the quality of DNA samples. Size selection (Blue Pippin, Sage Science, Beverly, MA, USA) was used to exclude DNA fragments between 10-250bp. The final concentration of DNA within the 250-1200 bp range was 5.213 nM, eluted to approximately 30 µl. The average fragment length was 434 bp. Pooled DNA was sequenced using the Illumina Miseq Standard v.3 2×300bp sequencing platform at the Oregon State University Core facility (Corvallis, OR, USA).

In addition to our samples, we included a pure-water negative control and two positive mock community controls in order to address any downstream sequencing or bioinformatic inconsistencies. Our mock community was created in conjunction with a shared study, see Thomas et al (2019) for details. Taxonomic identities of the mock community are shown in supplementary materials (Supplementary Table 2).

### Bioinformatics

The bioinformatics pipeline was based on the USEARCH/UPARSE pipeline (Edgar 2013) and follows Thomas et al. (2019). Reads were trimmed to remove low quality base calls and chimeric sequences, using a phred cutoff score of 34. Forward reads were trimmed to 250 bp, while reverse reads were trimmed to 220 bp. After quality checks, OTUs were generated using a 97% similarity cutoff. Taxonomy was assigned to fungal OTUs by BLASTing against a curated database, UNITE ITS (Abarenkov et al. 2010, Koljalg et al. 2013). We used variance stabilization as a method to deal with sequencing depth biases using the DESeq2 package in R (Love et al. 2014). Positive and negative controls were subtracted from all observations to reduce error from index-misassignment. Fungal ecological guilds were assigned using the FUNGuild tool introduced by Nguyen et al. (Nguyen et al. 2016). Full scripts of the data and bioinformatic pipeline are available as supplementary information.

### Statistical analyses

Because read abundance doesn’t necessarily translate to copy number (Nguyen et al. 2015), all analyses were performed on incidence (presence/absence) data. We first examined composition and diversity of both full fungal communities and ecological guilds of both host species. We then looked at full community and guild structure and at what the major drivers are that structure these communities.

All statistical analyses were performed using the R statistical environment (version 3.5.2, (R Core Team 2018)) using the vegan (Oksanen et al. 2017) and phyloseq (McMurdie and Holmes 2013) packages. Figures were created using the ggplot2 package (Wickham 2016). All tests requiring the use of dissimilarity matrices were performed using Jaccard distances (Oksanen et al. 2017).

#### Community composition

We calculated OTU richness for both full community and ecological guilds

#### community structuring

We used permutational multivariate analysis of variance (PERMANOVA) to model fungal community dissimilarity among and between host, ecoregion, and site. We used a permutation-based test of multivariate homogeneity of group dispersions to verify PERMANOVA assumptions (Veach et al. 2016). To visualize differences in fungal community composition we used non-metric multidimensional scaling (NMDS).

#### spatial structuring

To examine linear spatial structuring and spatial autocorrelation of fungal communities we used multivariate Mantel tests to test the correlation between our geographic distance matrix and our community dissimilarity matrices. Given the large geographic distance separating sites, our geographic distance matrix was computed using haversine distances. Mantel tests were visualized using Mantel correlograms. However, because we wanted to test the contribution of spatial distance on fungal community composition, we modeled spatial eigenvectors using principal coordinates of neighbor matrices, or distance-based Moran’s eigenvector map analysis (PCNM/dbMEM). These vectors were created following (Borcard et al. 1992). Briefly, we calculated a distance matrix from latitude/longitude values and used principal coordinates analysis (PCoA) to produce eigenvectors. Those with positive values were regressed using redundancy analysis (RDA) against the community matrix (Hellenger transformed, (Legendre and Gallagher 2001)). We used forward stepwise model selection to filter out non-informative vectors.

#### partitioning of variation

We used variation partitioning to quantify the relative contribution of host fitness, environment, and spatial structuring on fungal community composition (Table 1). We modeled host fitness using host demographic measures of size and reproductive output, host density, and foliar damage from natural enemies (pathogens, herbivores, etc.). Environmental data was represented as the previous year’s data obtained from the Parameter elevation Regression on Independent Slopes Model (PRISM) 800m resolution dataset (PRISM Climate Group, Oregon State University). We transformed these raw environmental data into bioclimate predictors to capture seasonal mean climate conditions (temperature, precipitation, and dewpoint seasonality) and intra-annual seasonality (i.e., precipitation of wettest quarter, temperature of coldest quarter, etc.), (O’Donnell and Ignizio 2012). We also included elevation as a climate predictor because elevational changes can mirror those of latitude. Our positive eigenvectors from the PCNM analysis were used as spatial predictors. Variation partitioning has been developed as a technique aimed at establishing relationships among community data and environmental predictors and has the advantage that up to four explanatory matrices can be tested against a community response matrix. It does this through the use of direct gradient analysis (in this case RDA) to test the partial, linear effect of each explanatory matrix on the response data (Borcard et al. 1992). Despite the robust nature of this technique, the results are still sensitive to multicollinearity, which can lead to distorted results (Buttigieg and Ramette 2014). We used stepwise model selection to filter out highly multicollinear variables.

We created matrices for each major filter category (environment, host fitness, spatial structuring), and regressed them against our community matrix. We used the function *varpar()* to perform the analysis. This function gives an output listing adjusted R^2^ values for all combinations of explanatory matrices, including host fitness (a), spatial distance (b) and abiotic (c), as well as shared partitioning for host and spatial (a & b), environment and spatial (b & c), etc. We used db-RDA to test each individual fraction, and used Monte Carlo global permutation tests of significance of canonical axes. We plotted the results to visualize the contribution of each fraction on endophyte community composition.

Finally, after partitioning the variation with respect to the full community, we used Funguild to breakout the fungi for which functional attributes are known into three functional groups (saprotrophs, pathotrophs and symbiotrophs) (Nguyen et al. 2016). Our initial pool of OTUs were reduced because we filtered out members which were either unassigned or low-confidence guild assignments. We used type III ANOVAS to examine ecological guild richness among sites. We partitioned variation for each ecological guild group as described above.

## Results

### Fungal Diversity and Composition

A total of 11,033,436 raw sequences were returned from our Illumina MiSeq run. After extensive quality control 9,432,696 sequences remained, 1,182,213 of which were unique. Following our clustering steps and further quality filtering, we detected a total of 3713 OTUs. Of those, *F. roemeri* was host to 1411 unique OTUs, while *D. californica* was host to 887 OTUs. These hosts shared a total of 1415 OTUs (Supplemental Table 1).

The full fungal community was comprised of seven phyla, 27 classes, 103 orders, 220 families, and 599 genera. Despite significant differences in fungal richness (Fig. S4), and taxa (Fig. 1 and 2, Supplemental Table 1), the phylum and class-scale fungal taxonomic diversity was relatively similar among host species. Within both *F. roemeri* and *D. californica*, the two most well represented phyla were Ascomycota (Fig. S4). For both host species, the five most common classes were Agaricomycetes, Dothideomycetes, Eurotiomycetes, Sordariomycetes, and Tremellomycetes (Fig. S5). We found that richness of communities of *F. roemeri* depended on site (F_6,77_ = 2.94, p = 0.012, Fig. S4a), but there was not a strong latitude component; instead French Flat had the lowest and nearby Table Rock was the highest. We found no difference in richness among sites for communities of *D. californica* (F_5,65_ = 1.84, p = 0.1173, Fig. S4b).

**Figure 1.**
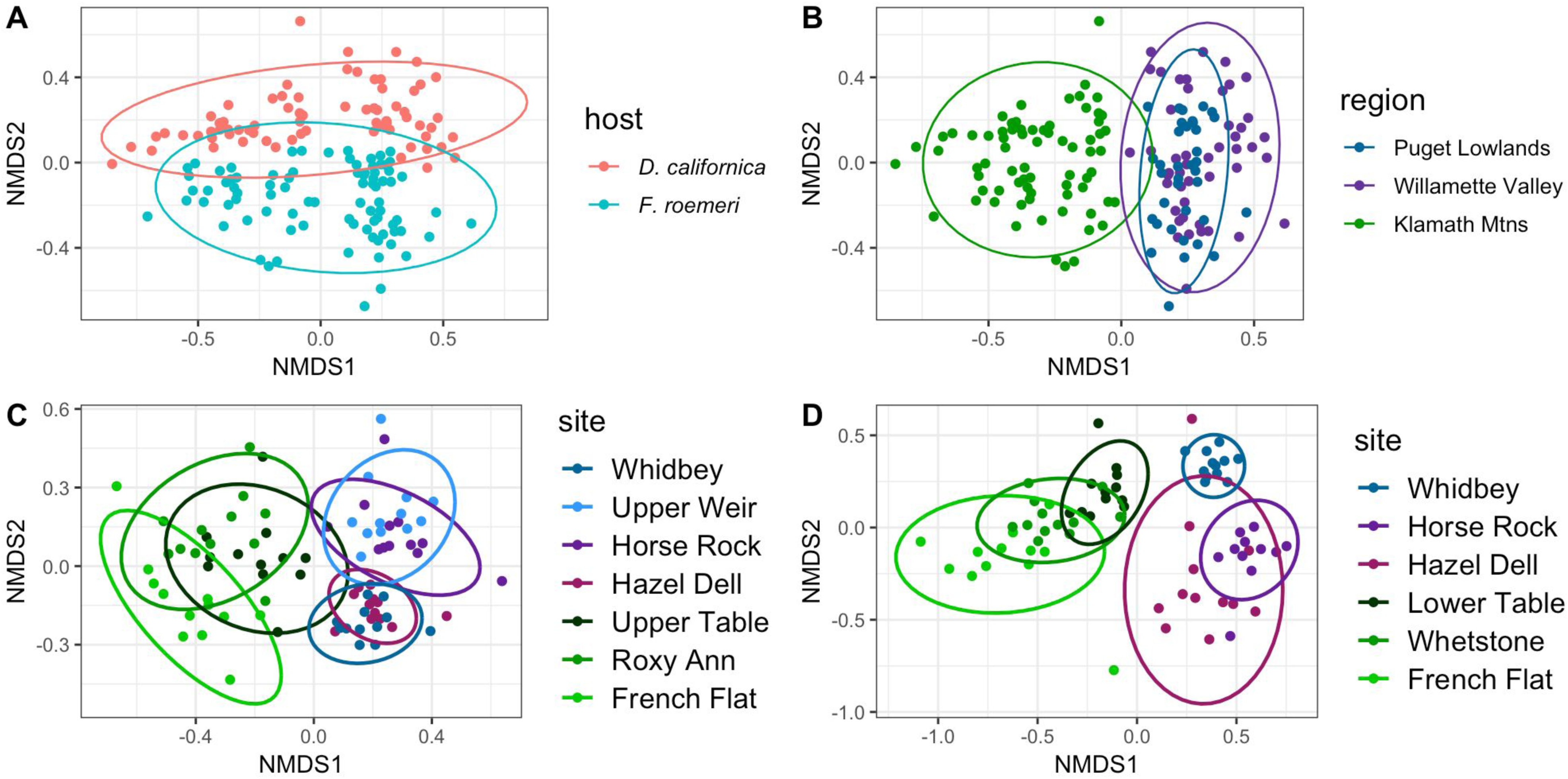
Non-metric multidimensional scaling (NMDS) plots showing relationship of fungal communities to (A) host species, (B) Ecoregion, (C) *F. roemeri sites*, and (D) *D. californica* sites. Ellipses represent 95% confidence intervals.

**Figure 2.**
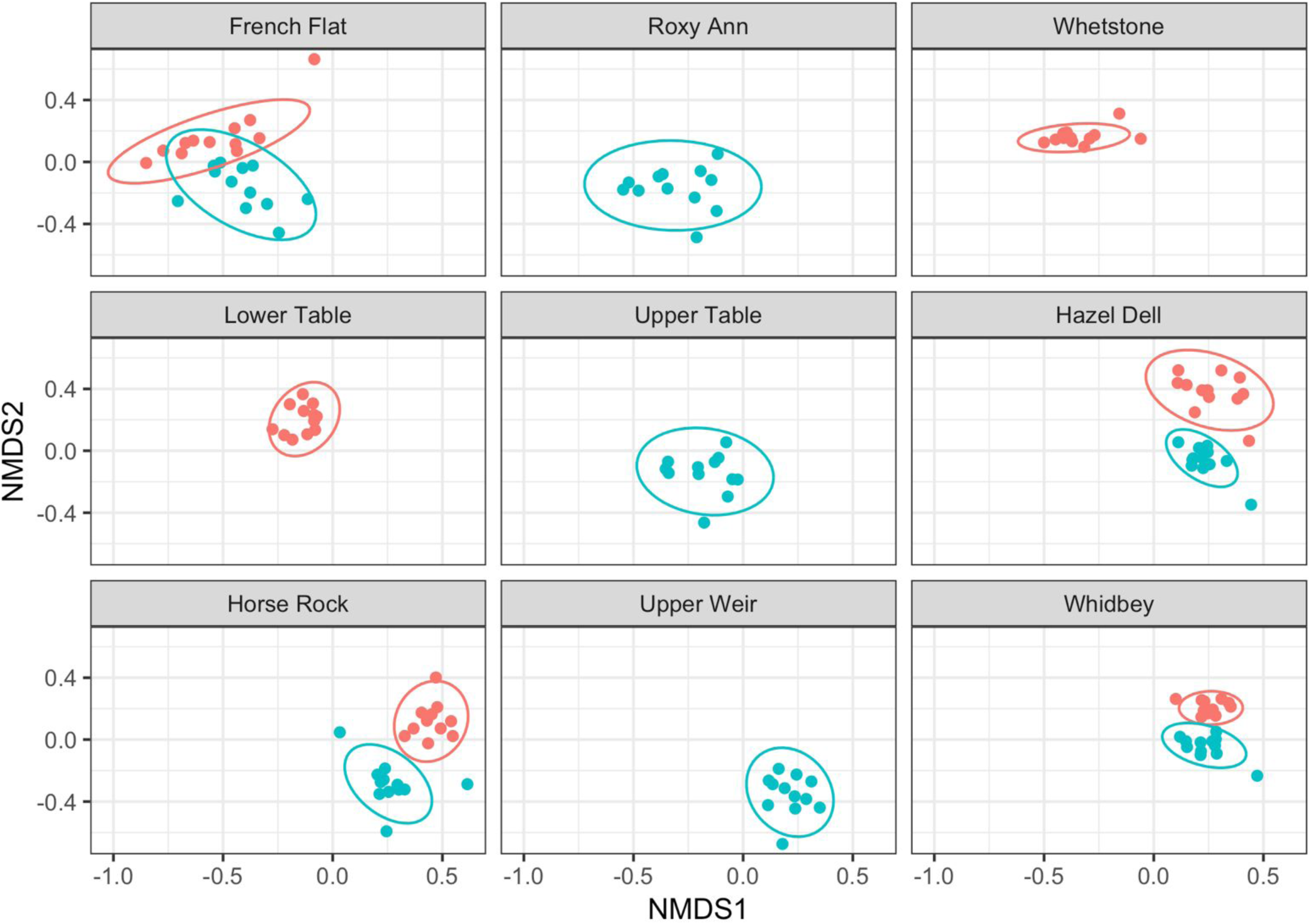
Multi-panel non-metric multidimensional scaling plots showing community structure separated by host at each site.

Of our initial 3713 OTUs, 869 were unassigned to an ecological guild. After we filtered out any low-confidence guild assigned OTUs (confidence ranking of ‘possible’), we were left with 2060 OTUs, 1577 associated with *F. roemeri*, and 1262 associated with *D. californica*. The common fungal classes remained the same as the full community. Communities of *F. roemeri* tend to increase in symbiotroph (F_6,77_ = 6.98, p < 0.001, Fig. 5a) and pathogen richness (F_6,77_ = 5.04, p < 0.001, Fig. 5b) with increasing latitude. However, we found that saprotroph richness was not different among sites (F_6,77_ = 2.07, p = 0.07, Fig. 15c). The final category is for the fungi that are not known sufficiently to fit into one of the trophic categories (uncategorized) and those that are unidentified to species; we include those in the final figure because they are part of the uncovered biodiversity and they too vary by site (F_6,77_ = 2.44, p = 0.03257, Fig. 5d). For *D. californica*, there was a slight tendency for symbiotroph (F_6,77_ = 4.89, p < 0.001, Fig. 6a) and pathogen richness (F_6,77_ = 7.44, p < 0.001, Fig. 6b) to increase with latitude. However, we saw decreasing richness of saprotrophs (F_6,77_ =11.93, p < 0.001, Fig. 6c) and unassigned/low confidence guild designations (F_6,77_ =5.16, p < 0.001, Fig. 2d) with increasing latitude.

**Figure 3.**
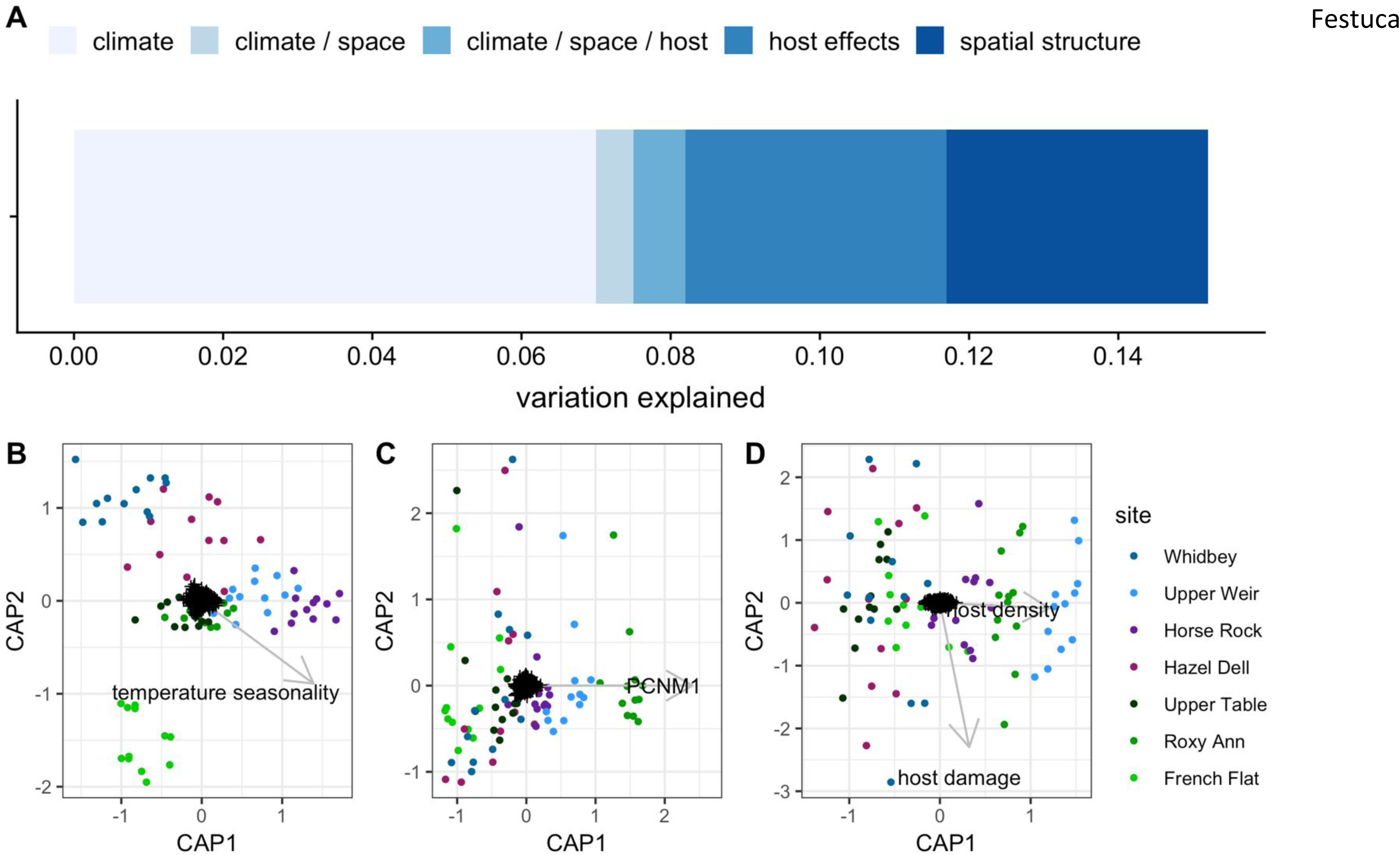
*Festuca roemeri*, RDA biplots showing the contribution of specific variables and total variance explained. (A) variance explained by each individual variable group (B) Biplot of Climate, (C) Biplot of spatial distance, (D) Biplot of host effects.

**Figure 4.**
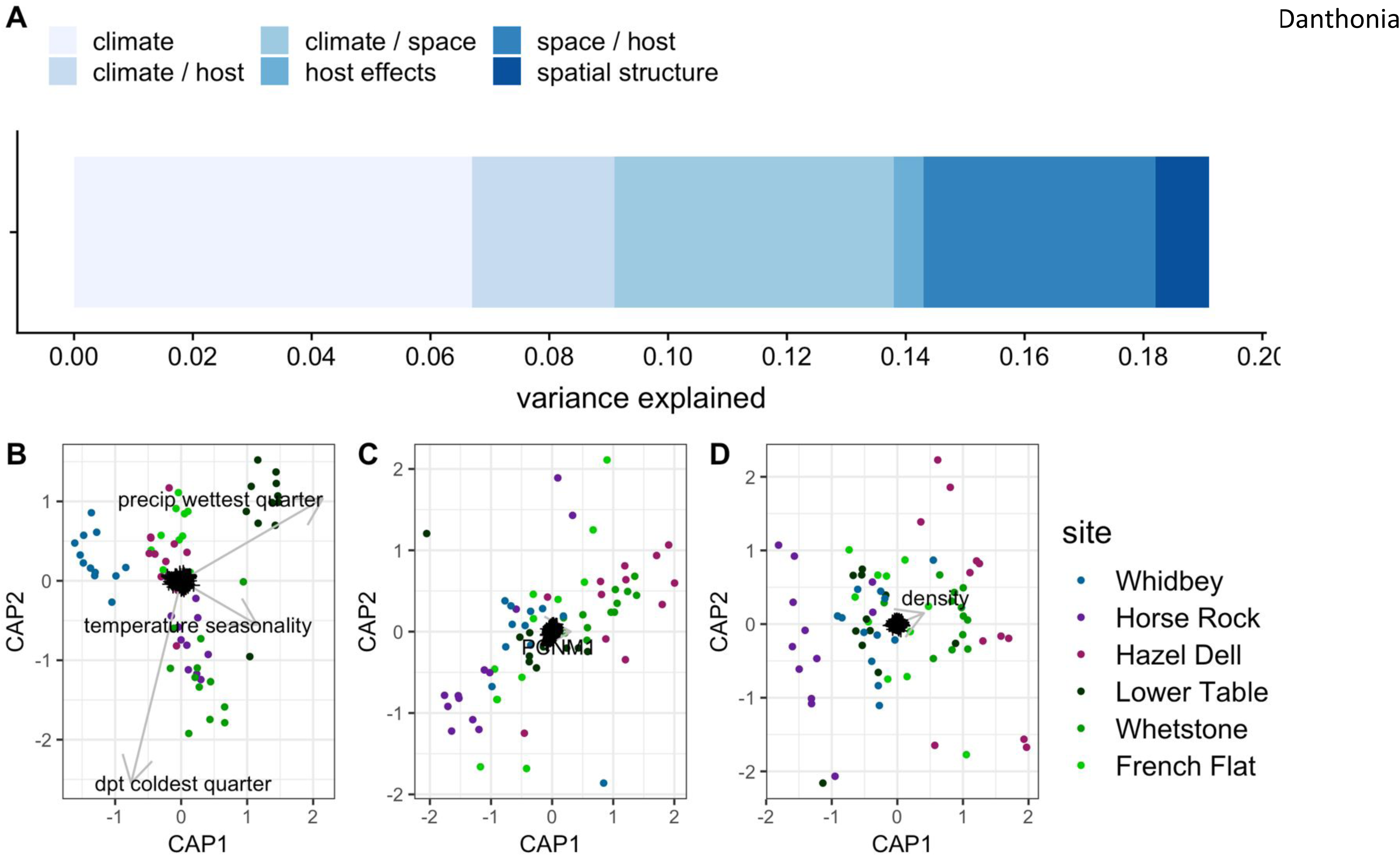
*Danthonia californica*, RDA biplots showing the contribution of specific variables and total variance explained. (A) variance explained by each individual variable group (B) Biplot of Climate, (C) Biplot of spatial distance, (D) Biplot of host effects.

**Figure 5.**
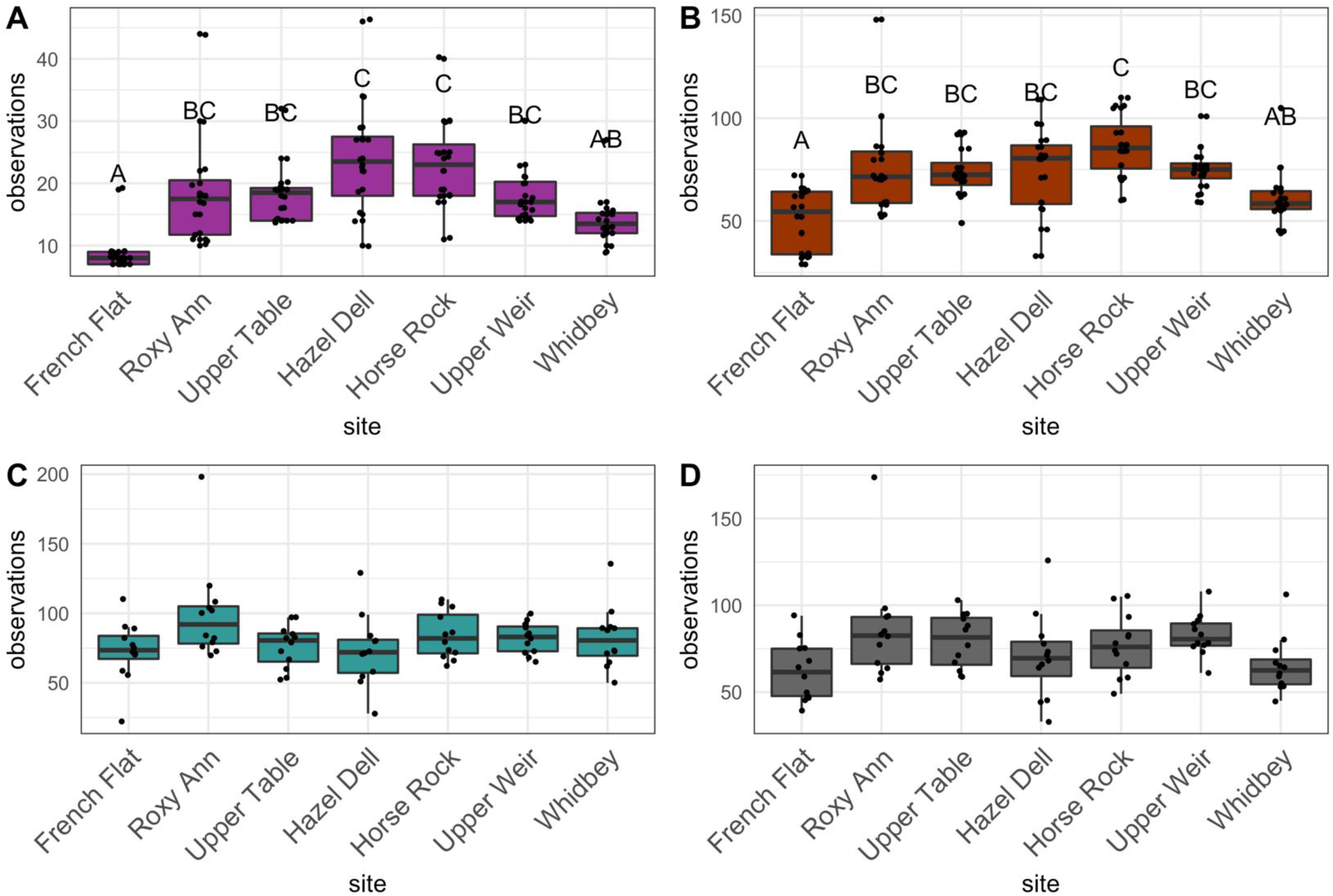
Guild richness for *Festuca roemeri*. (A) Symbiotrophs, (B) Pathotrophs, (C) Saprotrophs (D) Unassgined and low-confidence guild assignments. Sites are ordered by latitude.

**Figure 6.**
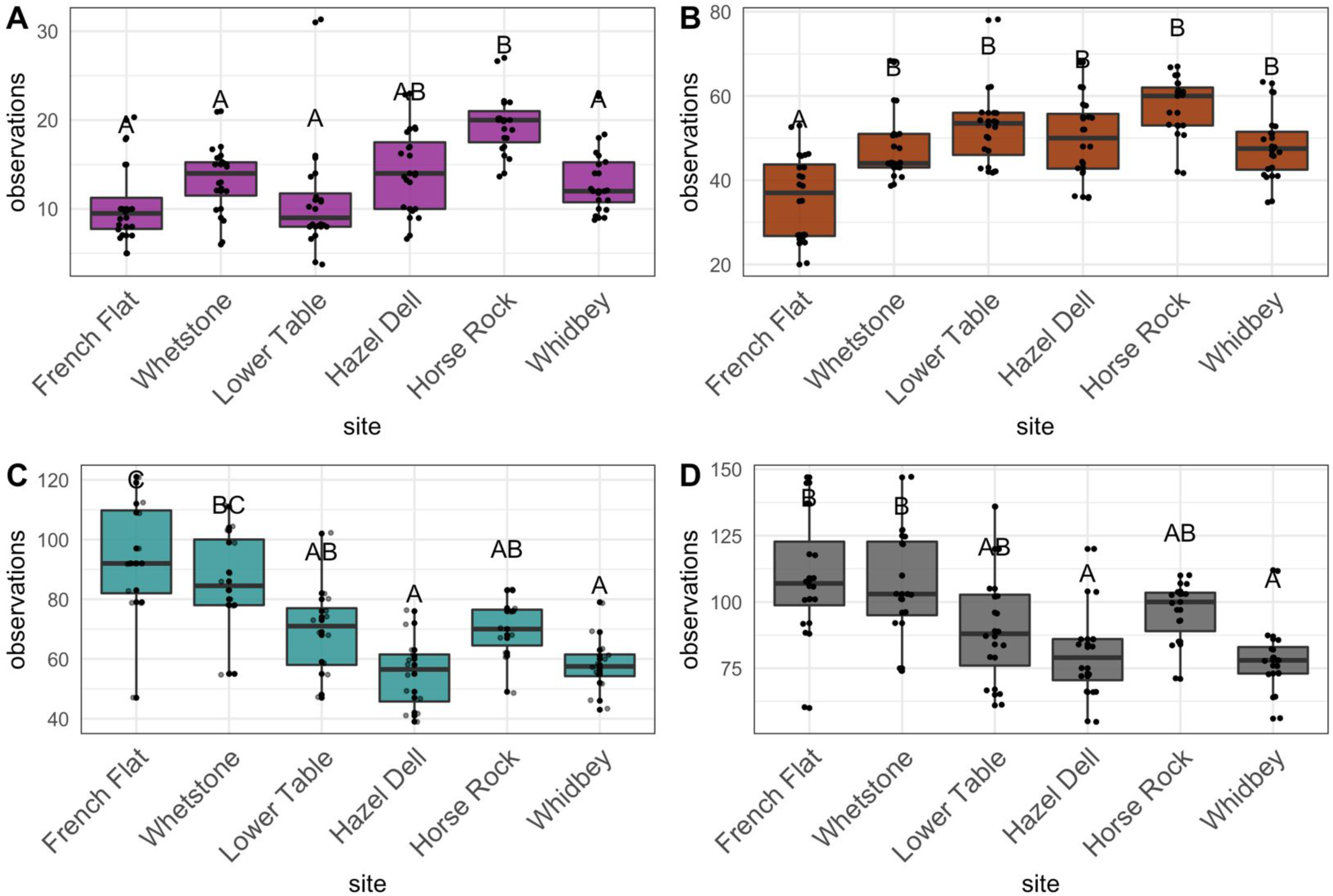
Guild richness for *Danthonia californica*. (A) Symbiotrophs, (B) Pathotrophs, (C) Saprotrophs (D) Unassgined and low-confidence guild assignments. Sites are ordered by latitude.

### Fungal community structure

In the full model, which included both host species, we found that host is a significant yet weak predictor of total fungal endophyte community structure (F_1,153_ = 5.67, p < 0.001, R^2^ = 0.036, Fig. 1a). Interestingly, when both hosts were present at the same site, their communities showed little to no overlap (Fig. 2). Ecoregion was also a relatively weak predictor for total fungi (F_3,153_ = 5.17, p < 0.001, R^2^ = 0.064); the southern sites formed a distinct cluster, while the central and northern sites clustered together (Fig. 1b). However, when the analyses are separated by species, site was a strong predictor variable for both *F. roemeri* (F_5,70_ = 3.22, p < 0.001, R^2^ = 0.20) and *D. californica* (F_5,70_ = 4.67, p < 00.1, R^2^ = 0.26). For *F. roemeri,* see distinct clustering separating sites from our southern ecoregion from those in the northern two (Fig. 1c). In contrast with those from *F. roemeri*, the *D. californica* communities showed stronger clustering by latitude. Communities clustered by region along the first NMDS axis, but also by latitude within those groupings along the second NMDS axis (Fig. 1d).

We found a weak pattern of community turnover in *F. roemeri* communities (Mantel’s r = 0.16, p < 0.001). Communities of *D. californica* showed strong patterns of community turnover (Mantel’s r = 0.45, p < 0.001). Within both host species, we saw positive autocorrelation between distance and community at closer distances (*F. roemeri* < 75km, *D. californica* < 175km), changing to negative spatial autocorrelation as distance increased (Fig. S3).

Given the large distances of our sampling scheme, we were only able to pick up on broad-scale spatial structures. For communities of *F. roemeri*, we found three positive PCNM vectors, all three of which were significant significant (PCNM1 F_1,80_=1.6002, p = 0.006, PCNM2 F_1,80_=3.0612, p < 0.001, PCNM3 F_1,80_=2.5417, p < 0.001). PCNM1 indicates a spatial structure in which greater similarity exists between the Klamath and Puget Trough regions, while the PCNM2 shows structuring which separates Klamath from both the Willamette Valley and Puget Trough, mirroring latitude. Similarly, for *D. californica* communities, we found three PCNM variables, two of which were significant (PCNM1 F_1,67_ = 5.0190, p < 0.001; PCNM2 F _1,67_ = 1.2010, p = 0.074, PCNM2 F _1,67_ = 6.1488, p < 0.001). Again, both PCNM variables map to broad scale spatial structures. PCNM1 seems to reflect the same split between Klamath and Willamette Valley/Puget Trough regions. PCNM2 again suggests a similarity between the Klamath and Puget Trough regions.

### Drivers of endophyte structuring

After stepwise selection, our final model for *F. roemeri* community structuring included four climate predictors (MAT, precipitation seasonality, temperature seasonality, and precipitation of the wettest quarter), one spatial predictor (PCNM1), and two host predictors (host foliar damage and host density). Our model for communities of *D. californica* was similar with four climate predictors (dewpoint temperature of the coldest quarter, temperature seasonality, and precipitation of the wettest quarter), one spatial predictor (PCNM1), and four host predictors (host size, host reproductive output, host foliar damage, and host density). The combined effect of climate, host and spatial-related variables explained 14.1% of the total community variation in *F. roemeri* (F_7,76_ = 2.9431, p < 0.001). Climate was the strongest predictor of community structure, explaining 7.8% of unique community variation (F_4,76_ = 2.8136, p < 0.001, Fig. 3a), followed by spatial structuring (4.1%, F_1,82_ = 4.6895, p < 0.001) (Fig. 3a). However, host fitness explained very little variation (4.2%, F_2,81_ = 2.945, p < 0.001) (Fig. 3a). The contribution of individual variables are show in Figure 3(B-D) and table 2. The total variation explained for *D. californica* endophyte community structuring was 19.2% (F_8,62_ = 3.0735, p < 0.001). Similarly, climate explained 6.7% of the unique variation in *D. californica* endophyte community structure (F_3,62_ = 2.7846, p = 0.001, Fig. 4a). Host fitness explained 0.5% of unique community variation (F_4,62_ = 1.0967, p = 0.045, Fig. 4a), and spatial-structure explained 0.92% (F_1,62_ = 1.7138 p < 0.001, Fig. 4a). Individual variable contribution is shown in table 2 and Figure 4 (B-D). We observed the same general pattern of variable importance in each ecological guild, although the variation explained did vary (Table S2-S4).

**Table 2.** Results of full community partitioning of variation. Main component contribution (adjusted R^2^) is shown, along with the contribution of each individual variable (pseudo F-value). Significant p values are shown in bold.

| Full community |  |  |  |  |  |  |  |
| --- | --- | --- | --- | --- | --- | --- | --- |
| component | variable | F. roemerii |  |  | D. californica |  |  |
|  |  | adjusted R<sup>2</sup> | pseudo F | p value | adjusted R<sup>2</sup> | pseudo F | p value |
| Climate | mean dewpoint temp coldest quarter | 0.067 | - | - | 0.185 | 4.007 | **0.001** |
|  | temperature seasonality |  | 1.591 | **0.009** |  | 3.490 | **0.001** |
|  | precip wettest quarter |  | 2.439 | **0.001** |  | 3.934 | **0.001** |
|  | precipitation seasonality |  | 2.433 | **0.001** |  | - | - |
|  | mean annual temperature (MAT) |  | 1.203 | 0.093 |  | - | - |
|  | elevation |  | - | - |  | 3.861 | **0.001** |
| spatial structure | PCNM 1 | 0.052 | 5.527 | **0.001** | 0.096 | 8.435 | **0.001** |
| Host fitness | host size | 0.015 | - | - | 0.068 | 0.742 | 0.972 |
|  | host reproduction |  | - | - |  | 1.902 | **0.003** |
|  | host foliar damage |  | 1.330 | **0.041** |  | 1.049 | 0.333 |
|  | host density |  | 1.848 | **0.002** |  | 3.874 | **0.001** |

## Discussion

### Fungal community structure

It is clear that the foliar fungal endophytes examined in this study display a non-random distribution across Pacific Northwest prairies. Our results indicate that host species, host ecoregion, and site all contributed to structure the fungal communities. Host specificity is one of our clearest results, yet it is contradictory to much of the literature on grass endophyte assembly (Sanchez Marquez et al. 2012, Higgins et al. 2014). The establishment and maintenance of host specificity is likely a product of many factors, including host traits, host phylogenetic relatedness, and host range (Kembel and Mueller 2014). Host traits encompass a wide range of attributes, including leaf physiology, leaf chemistry, and life histories. While we measured neither leaf chemistry nor quantitative host life histories, there are differences in host traits between these species. *F. roemeri* leaves are very narrow and tough, likely making them more recalcitrant to decomposition, while those of *D. californica* are broad and relatively thin, likely making them easier to decompose. These traits could also affect the colonization ability and rate of growth of fungal endophytes and may explain why richness estimators predicted such a larger diversity of fungi within *D. californica* (Fig. S7). Another factor that could contribute to the levels of host-specificity observed is that the two host species are very distantly related. While both species belong to the family Poaceae, they are from very different lineages in the family. *D. californica* belongs to the PACMAD clade (Sanchez-Ken and Clark 2010), while *F. roemeri* belongs to the BOP clade (Clark et al. 1995, Soreng et al. 2015). There is evidence that phylogenetic distance may be just as important in determining community composition as environmental, spatial, or biotic interactions (Gilbert and Webb 2007, Kembel and Mueller 2014). While host specificity may act to confine endophytic range, it tends to lead toward closer co-evolutionary relationships. This has significant implications for pathogens, where specificity may allow the fungi to escape host defenses, and better exploit plant resources (Barrett and Heil 2012).

Climate was the single best explanatory component for the communities of both host species. We expected that seasonality would be a strong driver in community structure, but unfortunately for *D. californica*, precipitation seasonality was too strongly correlated with other variables to include in the analysis. Likewise, while precipitation seasonality was included in our model for *F. roemeri*, it did not play a role in explaining unique variation. However, temperature seasonality was an important climatic predictor for both host species. We chose to model climate as that data collected the year leading up to sample collection. While there is evidence that fine scale weather patterns are very important in determining pathogen transmission (Garrett et al. 2006), it is possible that we lost information on the long-term climate patterns of the area. Despite this, our study highlights the importance of using bioclimatic predictors to model endophytic fungal communities, while annual predictors such as MAT may be less informative. Given future predictions of increased climatic seasonality, fungal communities may be especially sensitive to these changes.

While spatial structure was clearly important, it provided relatively low explanatory power for community structure, and was hard to disentangle from the effects of climate and host, especially in communities of *D. californica*. For both host species, we found the highest degree of similarity in community composition among central and northern sites, which were relatively distinct from southern sites. This is not surprising considering the topography of the area. The southern sites are separated from the central and northern sites by the Klamath Mountains, which have an elevational extent from 244 to 2,134 m. Mountains represent a substantial dispersal barrier for fungi (Peay et al. 2010), and they could be acting to differentiate communities on several levels. Current endophyte distributions likely depend on both dispersal of fungi and host. Besides roadway openings, there is also little to no connectivity of grasslands between the Klamath Mountain Ecoregion and the Willamette Valley. This suggests that for the most part, any movement of endophytes must be mediated by wind, or another external force. Depending upon host specificity of fungal community members, differentials in dispersal abilities could lead to the geographic structuring we uncovered. On the other hand, the Columbia River acts as a barrier between the Willamette Valley and Puget Trough ecoregions. The river is likely a small barrier, and indeed, some scientists don’t consider them to be distinct ecoregions (Floberg et al. 2004). For *D. californica*, we were largely unable to disentangle the effects of dispersal limitation from environmental filters and host effects. Because of the relatively small number of intact, unrestored prairies, we were left with an uneven sampling scheme, where our spatial scale varied from the range of tens of meters to the range of hundreds of kilometers. Unfortunately, there are few to no prairies that could bridge these two scales. Although dbMEM modeling is robust to irregular sampling schemes (Blanchet et al. 2009), it relies on a minimum spanning distance that can effectively connect any two sampling locations. As previously mentioned, our sampling sites were separated by large distances, the largest being several hundred km. Consequently, we were only able to pick up on very broad scale spatial patterns. It is highly likely that finer-scale spatial structuring is important for endophyte community structuring.

While host fitness was the weakest predictor category, it is interesting to note that host density contributed most to this variation. Host density could be affecting endophyte communities in several ways. Given that we saw significant host specificity, host density could impact transmission rates of specialist endophytes. This would give further evidence of dispersal limitations, especially those we encountered via the Klamath mountains. Host traits, including host leaf chemistry and physiology have been shown to be important in determining endophyte community composition (Kembel and Mueller 2014). To our knowledge, this is the first study to attempt to link fungal endophyte community composition/structure with attack of host by natural enemies. While we were only able to score host plants for presence/absence of damage types, we expect that a more in-depth sampling (i.e. % of photosynthetic area removed) would yield stronger results.

### Fungal Diversity and Composition

As we expected, the phyllosphere of native Pacific Northwest bunchgrasses is a very diverse habitat. We found over 3000 distinct OTUs among 159 individuals of two grass hosts, and this is estimated to be on the low side of the actual fungal diversity within these plants. Indeed,, we found only about half of the estimated diversity (Fig. S7). While we sampled only 12 fewer individuals from *D. californica*, we found nearly 500 fewer OTUs than from *F. roemeri* communities. Interestingly, when we examined the higher-level taxonomy among these host species, there were few differences; the two host species had remarkably similar taxonomic composition at the phylum, class, and order levels, perhaps reflecting that both hosts were in the grass family, Poaceae; indeed at this higher taxonomy, our results were very similar to three other genera of grasses (Wemheuer et al. 2019). However, the actual species in each of our host species were markedly different (Fig. 1a and Fig. 2) and many OTUs/species were not shared. There was an overlap of 1415 OTUs among both hosts, yet 1411 OTUs were only observed in *F. roemeri*, and 887 were only observed in *D. calfornica*. Thus, a very large portion of the diversity in these grasslands lies hidden in the leaves of the plants. The degree of host specificity that we observed suggests that conservation of these hidden fungi will depend on saving their hosts (Blackwell and Vega 2018).

Interestingly, species richness did not follow a latitudinal gradient as we expected. For communities of *F. roemeri*, we found the lowest diversity at the furthest southern site (French Flat), while the highest diversity was shared between a different southern Oregon and Washington site (Upper Table and Upper Weir). These sites didn’t share ecoregion, similar precipitation, or temperature regimes. Within *D. californica,* there were no significant differences in diversity among sites, although there was a trend of increased diversity with increased latitude. It should be mentioned that French Flat was our only site where the plants were growing on serpentine soil. These soils are characterized by high concentrations of Ca and Mg, as well as trace metals, which are inhospitable to most plants (Brady et al. 2005). As such, there are often vastly different plant communities on these soils, with higher proportions of endemic species (Whittaker 1954). Because of this, we examined diversity with and without this site. Interestingly, when French Flat was not included in the analyses, we found no difference in diversity among sites. These findings contrast with other studies which found that species diversity increases as latitude nears the equator (Arnold and Lutzoni 2007). However, Arnold and Luzoni (2007) found that while there was higher species diversity in the tropics, they were dominated by only a few fungal classes, while the relatively small diversity of species in northern latitudes originated from more classes. One notable difference between our study and that by Arnold and Lutzoni is the local environment. Researchers often imagine that the environment becomes more ‘harsh’ as you move away from the poles. However, we found the opposite, that the most seasonal climate (which we consider to be more extreme) is at our furthest southern site, and this trend in seasonality decreases as latitude increases. Other differences that could contribute to this difference in latitudinal diversity include the limited latitudinal scope of the study (∼680 km), host species, and regional fungal species pools. In addition, while their study was culture based, ours was culture-independent. Studies with smaller scope in distance may be unable to pick up these large-scale diversity patterns (Altman 2011). In contrast, an endophytic lifestyle could be acting to shelter the symbiont from external pressures. Since internal tissues are shielded from desiccation (Thomas et al. 2016), they may insulate symbionts from the effects of latitude (Willig et al. 2003), inflating the importance of biotic interactions and dispersal limitation.

Interestingly, ecological guild richness had larger and varying differences in richness than the full community. In both *F. roemeri* and *D. californica*, we saw the greatest symbiotrophic richness in the Willamette Valley, with the lowest richness at the northern and southern of edge sites this study. A similar pattern occurred for pathogrophic richness, with the highest diversity observed at our mid-latitude sites. Other studies such as Wang et al (2019) found similar patterns, where pathogenic richness displayed a unimodal pattern, with the highest richness at their mid-latitude sites. Their study suggested that this shift was due to a change in subtropical and temperate bioms, our study region didn’t have such diverse climate. We predicted that more extreme environmental conditions would increase pathogen richness as a function of plant stress. However, these sites do not represent extremes in elevation, intra-annual seasonality, or mean seasonal climate conditions. In contrast, we found that saprotrophic richness in *D. californica* generally tracked with latitude, with the highest richness in our southern-most site. Despite finding no significant differences in richness among sites, we found contrasting patterns of richness in *D. californica* among ecological guilds. These results suggest that while total community structure.

### Species composition

There were some surprises in the species composition of the leaves (Supplemental Table 1). First, the “grass endophytes”, the often mutualistic C-endophytes, were rare. Of the 155 grass individuals sampled, only three individuals from three different sites contained a single known mutualistic species, *Neotyphodium uncinatum*. Furthermore, *N. uncinatum* only occurred in *D. californica* (3/71 individuals), which has only previously been observed in Festucoid grasses (Barker 2008). Thus, this is the first report of *Neotyphodium uncinatum* in *Danthonia*, a non-festucoid grass. There were also five species of *Acremonium* but none of them are currently known to be grass-symbionts; the genus *Acremonium* is polyphyletic, and most species, including the five we found, are no longer considered to be clavicipitalean (Summerbell et al. 2011). We mention one specifically, *A. asperulatum* because it was relatively common (26/84 *F. roemeri* and 24/71 *D. californica*), suggesting it may be playing a biological role in these grasses and warranting further study. Previous records of this species are from soils (Giraldo et al. 2012). We suspect that *Neotyphodium* and related clavicipitalean endophytes are more common in our host species than this study suggests because they are often highly localized in plant tissue such as leaf axils, stems, and meristems. They are rarely found in leaf tissue (which we sampled here), and their frequency can depend on season and age of the leaf (Clay 1994, Schardl et al. 2004).

Another surprise was the number of insect pathogens (e.g., *Ophiocordyceps heteropoda*, *Pochonia* and *Metarhizium*) and lichen species (e.g., *Usnea, Ramulina, Cladonia*) in the leaves. However, examination of the literature revealed that a wide range of fungi often occur as endophytes, including lichenicolous fungi (Huang et al. 2018, Moler and Aho 2018, Voglmayr et al. 2019) and insect pathogens (Razinger et al. 2018, Rondot and Reineke 2018). We expect that the kinds of fungi known to occur as endophytes will only increase as more NGS studies are published.

Finally, there were many macrofungi, including mushroom species, hiding in the leaves as endophytes, including: *Bovista plumbea, Lycoperdon spp, Coprinellus micaceus, Hypholoma fasciculare* and *Deconica (Psilocybe) montana*, all of which have also been detected in a study of above ground fruiting bodies in PNW prairies (Roy et. al. unpublished). One of the more interesting ascomycete endophytes we found was *Xylaria schweinitzii*, which is not known from fruiting bodies in the PNW. So, either this fungus is in the PNW and has not yet been found fruiting (unlikely as it is tropical), or it was a contaminant from the lab, or the sequence could have been from another fungus in the X. polymorpha clade, which have complex variations at the ITS locus and are not reliably identified with the ITS barcode (Roo Vandegrift pers. Comm.). We doubt that it is a contaminant because we were careful during sample preparation to use positive flow hoods, which were not in the lab where fungal specimens are observed, and the leaf surfaces were carefully surface sterilized. We think it is more likely to be a species in the *X. polymorpha* clade that is not identified well with ITS.

What are these fungi doing in the leaves? It is likely that in the wild many of these fungi find the moist interior of plant leaves to be a good place to “hide” from a drier external environment and some may also use the leaves for additional dispersal. Viaphytism is a recently coined term for this strategy (Nelson et al. in review, Thomas et al. submitted). Once thought to be largely confined to ascomycetes (Carroll 1999), NGS sequencing efforts are showing that hiding in plants is a common strategy for basidiomycetes too.

## Acknowledgements

This paper was derived from the master’s thesis of G. Bailes, which was completed under the guidance of B. Roy, with S. Bridgham, L. Pfeifer-Meister and B. Johnson as committee members. L. Hendricks, M. Krna, M. Petersen, W. Morris, and D. Doak aided with collecting field data; our apologies if we missed anyone who helped. Much appreciation goes out to all of the land managers/owners who allowed us to work on their land, including the Bureau of Land Management, the Center for Natural Lands Management, the City of Medford, Join Base Lewis McChord, the Nature Conservancy, the Oregon Department of Fish and Wildlife, and the Pacific Rim Institute. E. Alverson, R. Pelant, S. Hamman, S. Krock, C. Mayrsohn, M. Sullivan, and P. Dunwiddie all provided assistance in locating plant populations. D. Turnbull and M. Weitzman both provided advice and technical support in the preparation for, and the process of Illumina sequencing.

## Author Contributions

G. Bailes collected the data, did most of the analyses, and wrote the first draft of the manuscript. D. Thomas set up the mock communities with G. Bailes, and contributed greatly to the bioinformatic and statistical methodology used within this manuscript. S. Bridgham contributed to the conceptual development and editing. B. Roy contributed to the conceptual development, fieldwork, writing and editing.

## Supplemental Tables

Table S1: OTU table, available in final manuscript.

**Table S2.**
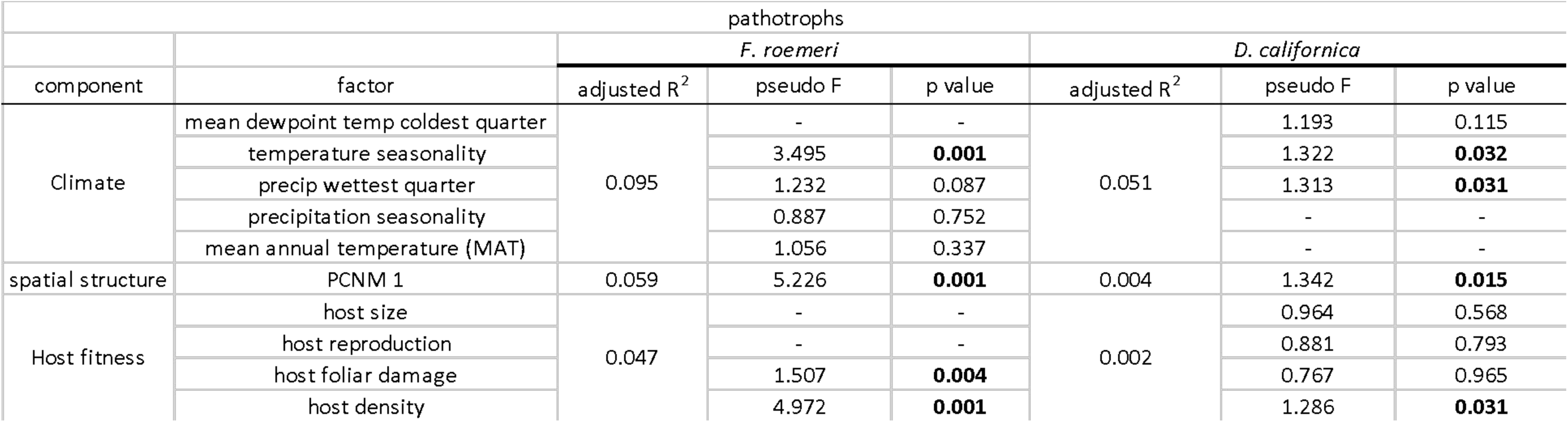
Results of pathogen partitioning of variation. Main component contribution (adjusted R^2^) is shown, along with the contribution of each individual variable (pseudo F-value). Significant p-values are shown in bold.

**Table S3.**
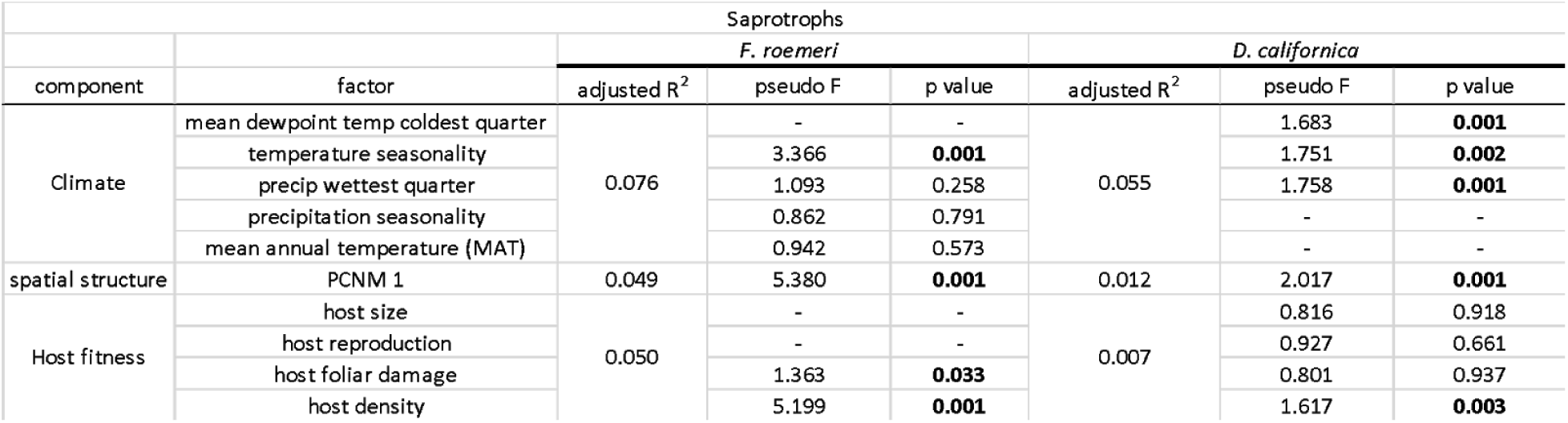
Results of saprotroph partitioning of variation. Main component contribution (adjusted R^2^) is shown, along with the contribution of each individual variable (pseudo F-value). Significant p-values are shown in bold.

| Saprotrophs |  |  |  |  |  |  |  |
| --- | --- | --- | --- | --- | --- | --- | --- |
| component | factor | *F. roemerii* |  |  | *D. californica* |  |  |
|  |  | adjusted R<sup>2</sup> | pseudo F | p value | adjusted R<sup>2</sup> | pseudo F | p value |
| Climate | mean dewpoint temp coldest quarter | 0.076 | - | - | 0.055 | 1.683 | **0.001** |
|  | temperature seasonality |  | 3.366 | **0.001** |  | 1.751 | **0.002** |
|  | precip wettest quarter |  | 1.093 | 0.258 |  | 1.758 | **0.001** |
|  | precipitation seasonality |  | 0.862 | 0.791 |  | - | - |
|  | mean annual temperature (MAT) |  | 0.942 | 0.573 |  | - | - |
| spatial structure | PCNM 1 | 0.049 | 5.380 | **0.001** | 0.012 | 2.017 | **0.001** |
| Host fitness | host size | 0.050 | - | - | 0.007 | 0.816 | 0.918 |
|  | host reproduction |  | - | - |  | 0.927 | 0.661 |
|  | host foliar damage |  | 1.363 | **0.033** |  | 0.801 | 0.937 |
|  | host density |  | 5.199 | **0.001** |  | 1.617 | **0.003** |

**Table S4.** Results of symbiotroph partitioning of variation. Main component contribution (adjusted R^2^) is shown, along with the contribution of each individual variable (pseudo F-value). Significant p-values are shown in bold.

| symbiotrophs |  |  |  |  |  |  |  |
| --- | --- | --- | --- | --- | --- | --- | --- |
| component | factor | *F. roemerii* |  |  | *D. californica* |  |  |
|  |  | adjusted R<sup>2</sup> | pseudo F | p value | adjusted R<sup>2</sup> | pseudo F | p value |
| Climate | mean dewpoint temp coldest quarter | 0.059 | - | - | 0.033 | 0.915 | 0.609 |
|  | temperature seasonality |  | 2.837 | **0.001** |  | 0.970 | 0.510 |
|  | precip wettest quarter |  | 1.186 | 0.155 |  | 1.031 | 0.403 |
|  | precipitation seasonality |  | 0.765 | 0.893 |  | - | - |
|  | mean annual temperature (MAT) |  | 1.004 | 0.448 |  | - | - |
| spatial structure | PCNM 1 | 0.027 | 3.345 | **0.001** | 0.000 | 0.987 | 0.509 |
| Host fitness | host size | 0.033 | - | - | 0.000 | 1.083 | 0.309 |
|  | host reproduction |  | - | - |  | 0.944 | 0.555 |
|  | host foliar damage |  | 0.907 | 0.647 |  | 0.691 | 0.965 |
|  | host density |  | 3.769 | **0.001** |  | 1.103 | 0.286 |

## Supplemental Figures

**Figure S1.**
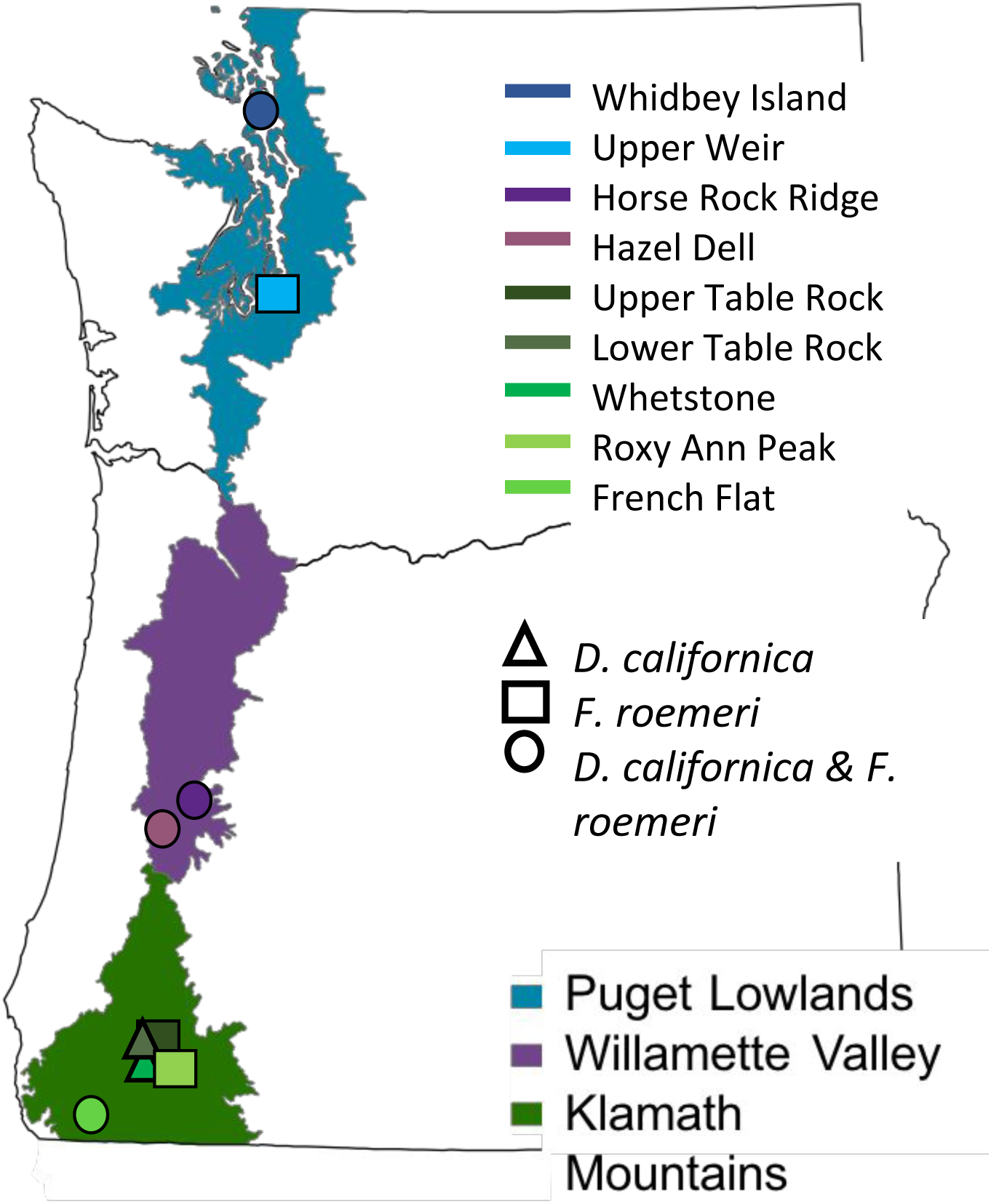
Map depicting ecoregions and area of study within Washington and Oregon. This Figure and all others following use the same color scheme: green represents the Klamath Mountain ecoregion, purple represents the Willamette Valley, and blue represents the Puget Lowlands.

**Figure S2.**
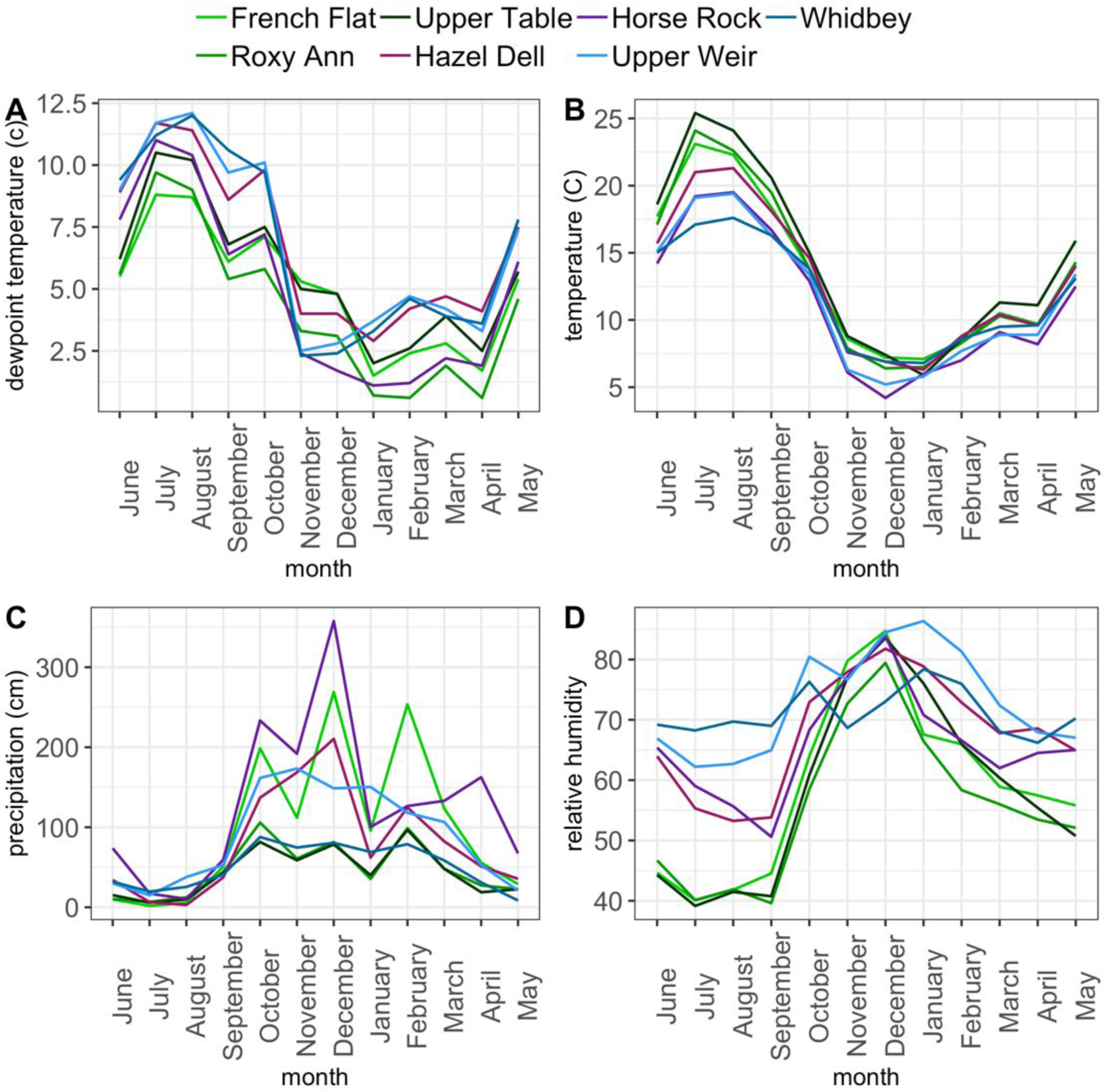
Mean monthly precipitation and temperature. Precipitation at (A) *F. roemeri* sites, (B) Precipitation at *D. californica* sites, (C) Precipitation at F. roemeri sites, (D) Precipitation at *D. californica* sites. Data are based upon 30-year normals (1980-2010).

**Figure S3.**
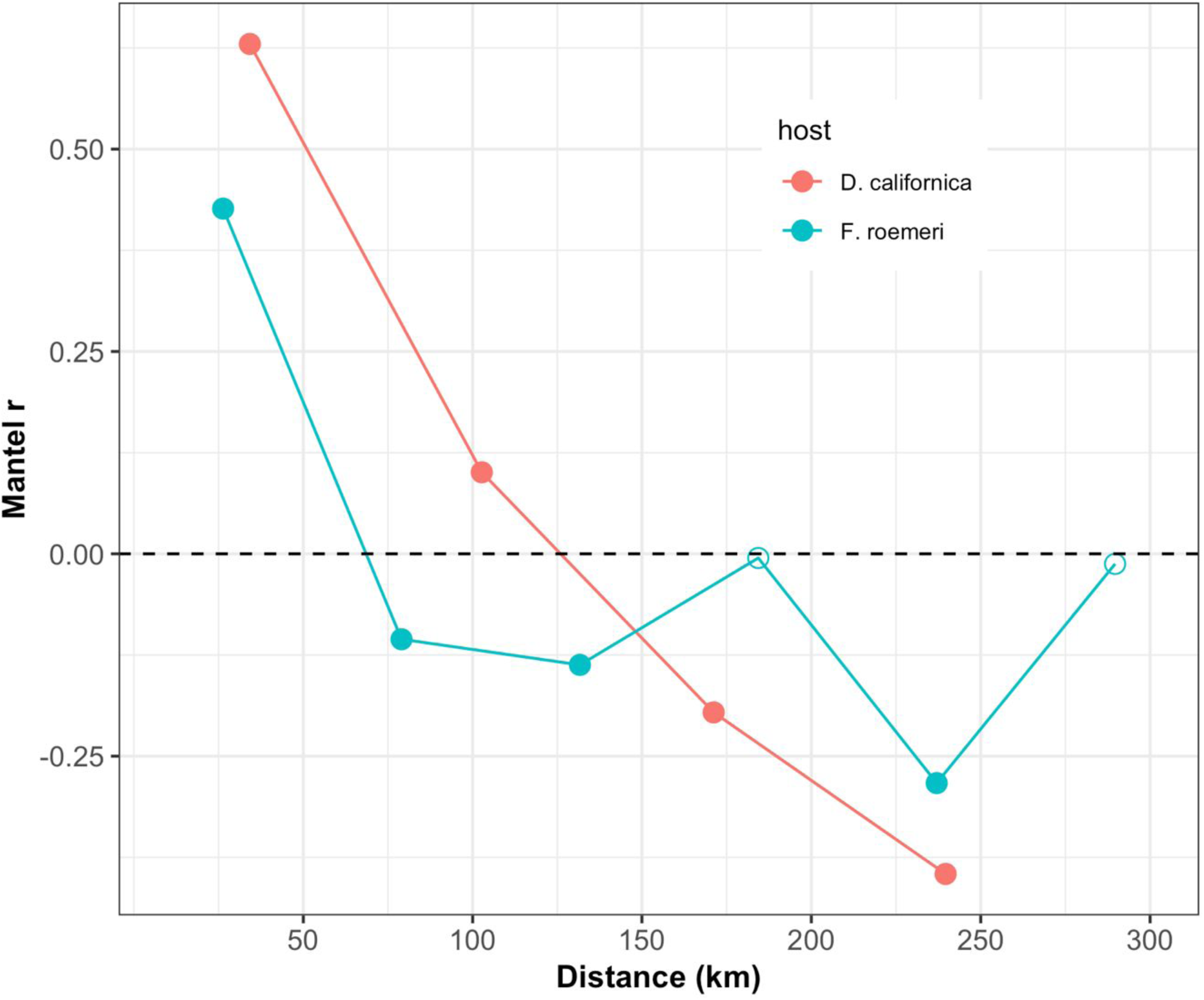
Mantel correlogram comparing community dissimilarity and distance among *F. roemeri* (Mantel’s r = 0.16, p < 0.001) and *D. californica* (Mantel’s r = 0.45, p < 0.001). Because a dissimilarity matrix was used, positive mantel’s r signifies spatial autocorrelation, while negative r signifies landscape homogenization. Filled circles represent significant comparisons.

**Figure S4.**
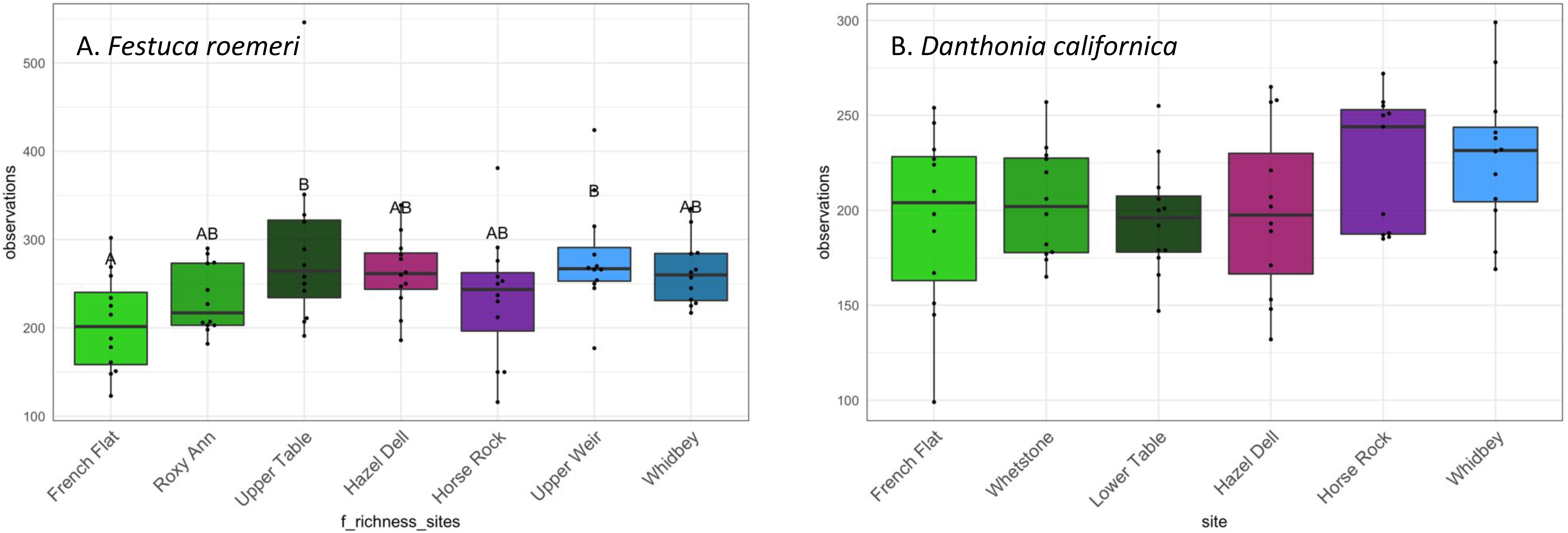
OTU richness by site for (A) *F. roemeri* and (B) *D. californica*. Shared letters are not significantly different by Tukey’s HSD. Error bars represent SE. Colors represent ecoregions: Klamath Mountains (green), Willamette Valley (purple), and Puget Trough (blue).

**Figure S5.**
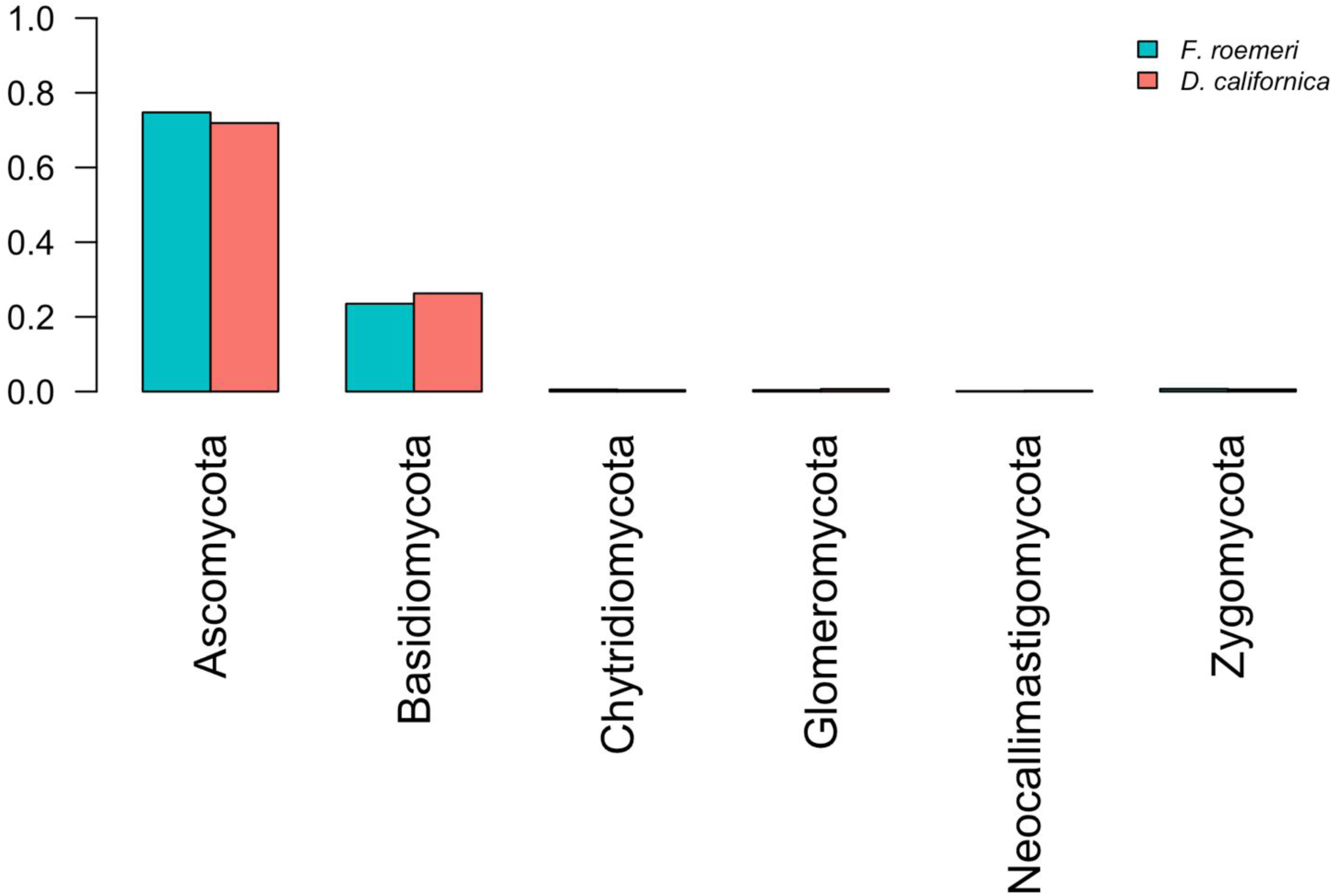
Major phyla recovered from both host species.

**Figure S6.**
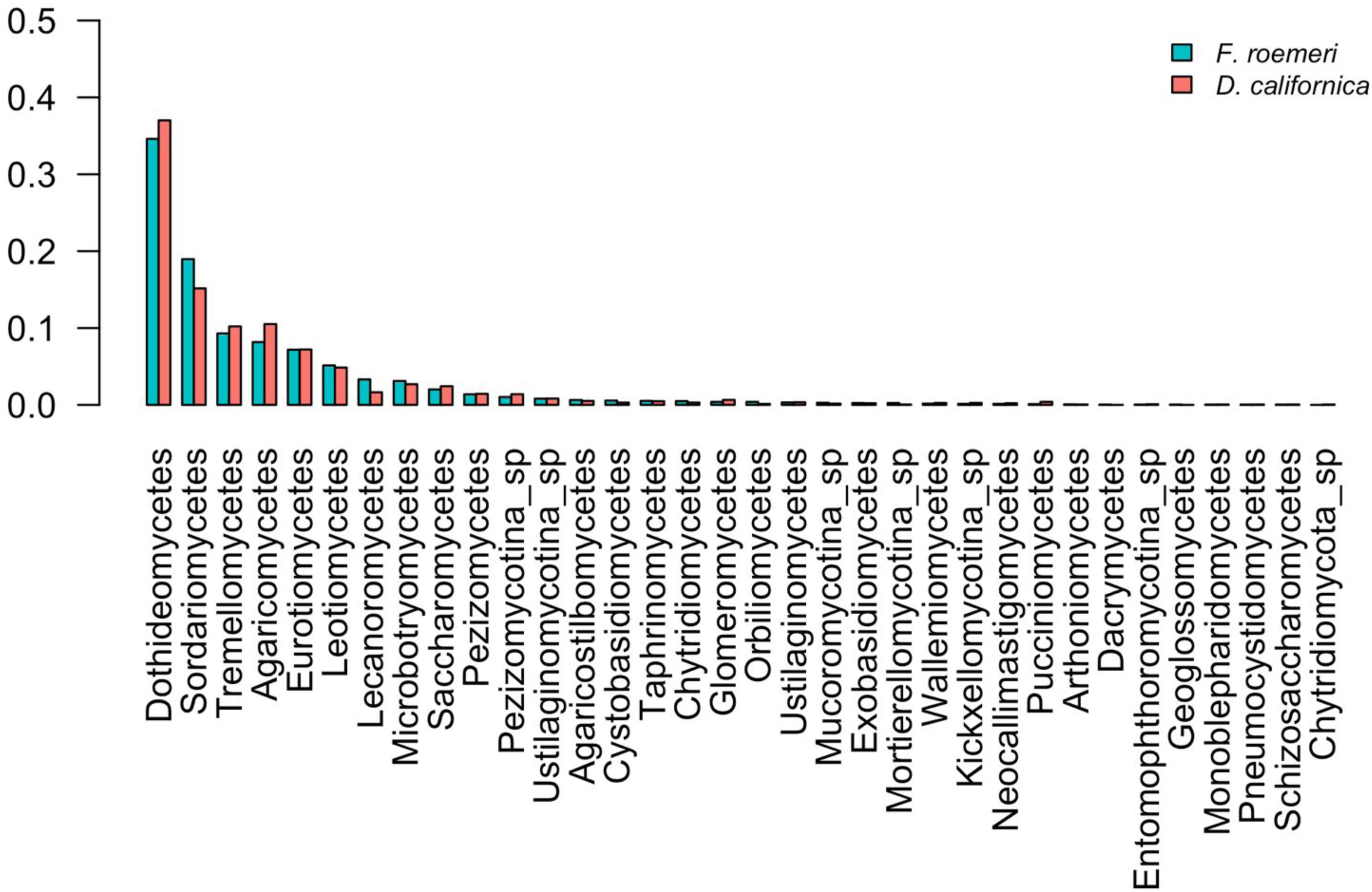
Major orders recovered from both host species.

**Figure S7.**
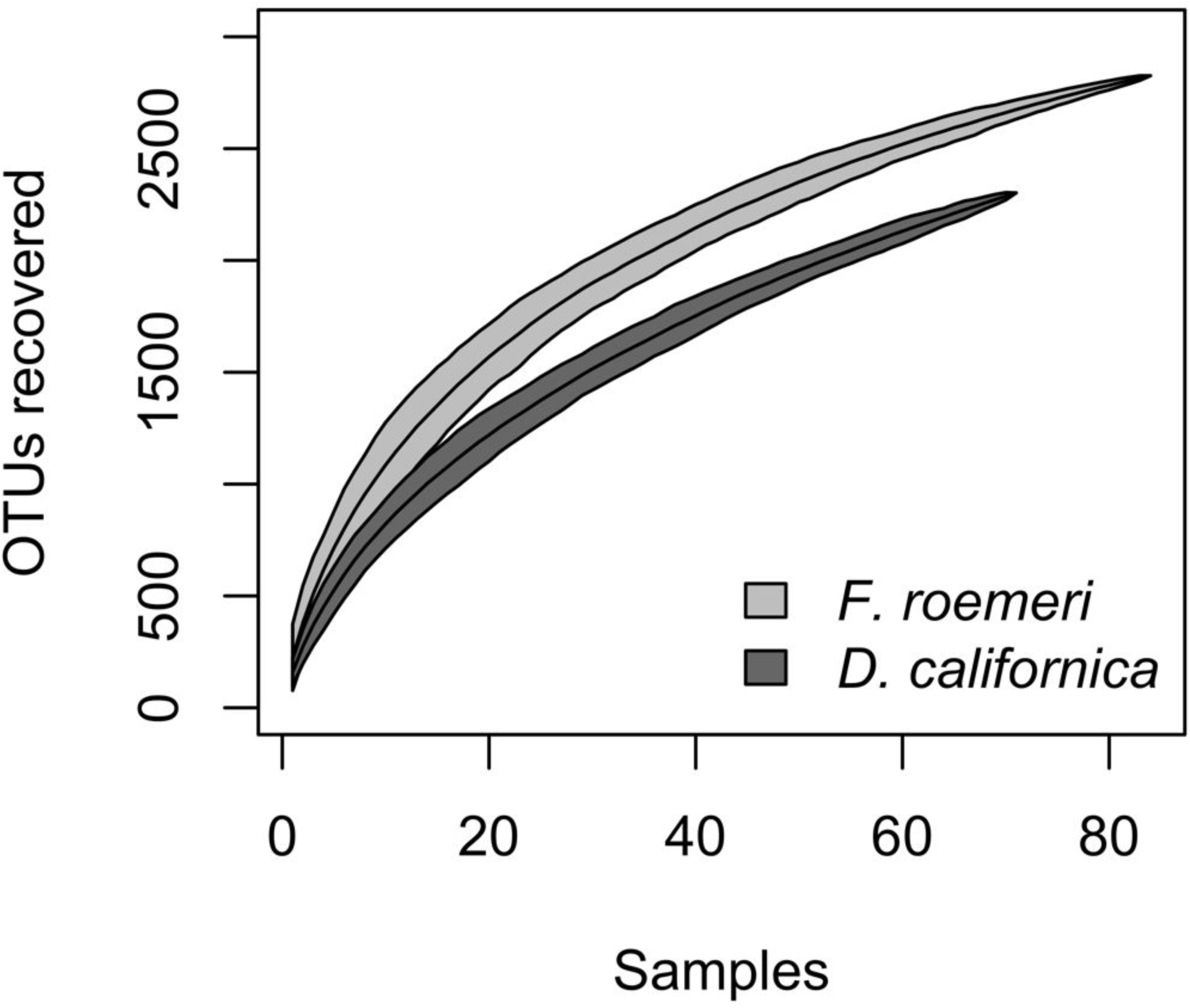
Fungal species accumulation curves for both host species.

